# *Drosophila beanbag* (*beba*) encodes a novel insect receptor tyrosine kinase associated with reproductive niche organisation

**DOI:** 10.64898/2026.07.06.736846

**Authors:** Sarah Mele, Sarah Bright, Emily Kerton, Travis K. Johnson

**Affiliations:** Department of Biochemistry and Chemistry, and La Trobe Institute for Molecular Sciences, La Trobe University, Bundoora, Victoria 3086, Australia

**Keywords:** Drosophila melanogaster, RTK, Ret, Torso, germline stem cell niche, hub cell, terminal filament, adult midgut progenitor, insect evolution

## Abstract

Receptor tyrosine kinases (RTKs) are cell surface proteins that govern many critical cell fate decisions and their dysregulation is a major cause of diseases such as cancer. Much of what we know about how these proteins work in cells and tissues comes from model organisms such as the fruit fly *Drosophila*. Here, we identify and characterise a previously unstudied *Drosophila* receptor tyrosine kinase encoded by CG3277, which we name *beanbag* (*beba*). Ectopic *beba* expression activated Akt and ERK phosphorylation and produced gain-of-function phenotypes resembling those caused by other *Drosophila* RTKs. Using a MiMIC-derived T2A-GAL4 allele, we show that *Drosophila beba* is expressed in digestive, nervous and reproductive systems, in locations suggestive of potential roles in endoreplication and/or stem cell niche support. Animals transheterozygous for *beba* loss-of-function alleles were viable, developed at a normal rate, and showed no detectable change in enterocyte DNA content under standard conditions. However, *beba* loss-of-function females had fewer ovarioles, consistent with a role in the ovarian terminal filament, and males had increased testis hub cell number and hub volume, suggesting *beba* may regulate somatic niche architecture in the *Drosophila* gonad. Phylogenetic analysis places Beba within a Ret/Tor-related RTK radiation and supports the existence of a distinct Beba family in insects. Together, our data define Beba as a lineage-restricted *Drosophila* RTK with specialised roles in reproductive niche organisation.

## Introduction

Receptor tyrosine kinases (RTKs) are transmembrane signalling proteins that couple extracellular cues to intracellular pathways critical for development and organismal homeostasis (Lemmon and Schlessinger 2010; Schlessinger 2014). Through these pathways, RTKs regulate diverse processes including cell and tissue patterning, cell shape and migration, cell and organ growth, and adult tissue maintenance and survival (Jin et al., 2015; Hu and Olsen, 2016; Boulan et al., 2019). RTK misregulation is a major driver of developmental disease and cancer, making them central to both basic biology and biomedical research. In animals, RTKs typically contain an extracellular ligand-binding region, a single-pass transmembrane domain and a cytoplasmic tyrosine kinase domain that activates downstream signalling after receptor dimerisation and phosphorylation (Lemmon and Schlessinger 2010; Schlessinger 2014).

Model organisms such as the mouse, zebrafish, fruit fly (*Drosophila melanogaster*), and the nematode worm (*Caenorhabditis elegans*) have provided deep insights into the roles, functions and regulatory mechanisms of RTKs and their signalling pathways, particularly in the contexts of animal development and disease (Shilo 2003; Lemmon and Schlessinger 2010; Schlessinger 2014). For example, studies of the insulin/IGF-like DAF-2 signalling pathway in *C. elegans* helped establish insulin/IGF-like signalling as a conserved regulator of lifespan and stress resistance (Kenyon et al. 1993; Morris et al. 1996; Kimura et al. 1997; Holzenberger et al. 2003). Similarly, studies of the EGFR signalling pathway in *Drosophila* identified conserved mechanisms of ligand processing and trafficking, as well as regulatory mechanisms including feedback loops and secreted ligand antagonists that influence signalling range (Schweitzer et al. 1995; Golembo et al. 1996; Urban et al. 2001; Shilo 2003).

*Drosophila* has been particularly valuable for understanding RTK biology because it combines genetic tractability, rapid life cycle and low cost with relatively low RTK family redundancy. The human genome encodes approximately 58 RTKs across 20 families (Manning et al. 2002; Lemmon and Schlessinger 2010), whereas the *Drosophila* genome encodes approximately 19 known RTKs, with single representatives for 11 of the 20 mammalian families (Mele and Johnson 2020; Gramates et al. 2022). This reduced complexity simplifies genetic analysis and avoids potential confounding receptor interactions, such as the heterodimeric relationships among vertebrate ErbB family members, for example (Yarden and Sliwkowski 2001). At the same time, the smaller *Drosophila* RTK complement includes receptors that are highly conserved with mammalian counterparts as well as receptors that are lineage-restricted or have no clear one-to-one vertebrate orthologue (*e.g.,* Torso and Sevenless; Nüsslein-Volhard et al. 1987; Sprenger et al. 1989; Casanova and Struhl 1989). This makes *Drosophila* useful not only for dissecting conserved RTK pathways, but also for understanding how RTKs diversify to control tissue-specific developmental processes.

Here, we report the identification of CG3277, which we name *beanbag* (*beba*); a previously uncharacterised gene with strong sequence similarity with the RTK Torso in its predicted tyrosine kinase domain. *beba* was initially annotated as a non-receptor tyrosine kinase because an expressed sequence tag aligned only with its tyrosine kinase domain-encoding region. Here, we provide evidence that *beba* encodes an RTK and further reveal its expression pattern in the digestive, nervous, and reproductive systems. Our functional studies implicate *beba* in the organisation of specialised niche support cells in male and female gonads, while phylogenetic analysis suggests that Beba represents a distinct insect-specific family of RTKs.

## Materials and Methods

### *Drosophila* stocks, generation and maintenance

The following stocks were obtained from the Bloomington Stock Center: UAS-*RedStinger* (BL8546), UAS-*mCD8-GFP* (BL5130), *beba^MI06697^* (BL43078), *w^1118^* (BL5905), *Df(2L)ED4651* referred to here as *beba^df^* (BL8904), *ptc*-GAL4 (BL2017), *actin*-GAL4 (BL25374), GMR-GAL4 (BL9146), and *en*-GAL4 (found in BL83350). The *beba^MI06697^* allele was converted to *beba^T2A^*^-*GAL4*^ via recombination-mediated cassette exchange using a previously described crossing scheme (Venken et al. 2011; Diao et al. 2015; Lee et al. 2018) using stocks BL60310, BL60299 and BL60291 for the Trojan exon donor, source of recombinase and integrase, and for screening recombinants, respectively. *beba* alleles were rebalanced over the CyO, GFP balancer (from BL9325). Multiple UAS-*beba* transgenic lines were created by gene synthesis of its open reading frame (Genscript) and insertion into pUASTattB *via* EcoRI/XbaI followed by transgenesis via phiC31 integration into the ZH-86Fb site (Bischof et al. 2007). Stocks were maintained on sugar-yeast food media at 22°C. All experiments were performed at 25°C.

### Immunoblotting

Three replicates of five flies overexpressing *beba* under the control of GMR-GAL4 (and GAL4 alone controls) were decapitated and their heads crushed in ice-cold lysis buffer (50mM Tris-HCl pH 7.5, 150mM NaCl, 2.5mM EDTA, 0.2% Triton-X, 5% glycerol) containing proteinase inhibitors (Roche). Protein lysates were normalised *via* BCA assay (Thermo Fisher Scientific) with 20 μg total protein separated via SDS-PAGE (Bio-Rad), blotted onto PVDF membrane which was then blocked with 5% skim milk powder in TBS-Tween (0.2% vol/vol), and probed with either anti-pAkt (Cell Signaling #4054, 1:1000) or anti-dpERK (Cell Signaling #4370, 1:1000) overnight in TBS-Tween containing 5% BSA. Blots were developed with ECL (Amersham) following secondary antibody incubation (anti-rabbit HRP 1:5,000, Southern Biotech) in block solution and further washing. Band intensities were quantified in ImageJ relative to prominent non-specific bands (Schneider et al. 2012).

### Quantitative gene expression analysis

RNA was isolated from groups of 10 whole larvae using Trizol (Thermo Fisher Scientific) and normalised *via* dilution before cDNA synthesis with oligo(dT) and random hexamers (Bioline). Quantitative PCR was performed on a LightCycler (Roche) with SYBR Green chemistry. Primers for *beba* targeting exons downstream of the trojan exon insertion (F-5’ TCC TTT GCC GAA ACC ACT TAC ACC 3’, R-5’ CTC TTC CAG GAA ACG CAT CCC A 3’) were designed (Primer3), validated and tested for efficiency. Reactions for *beba* and the control gene *Cyclin K* (Johnson et al. 2009) were run in triplicate for four biological replicates per genotype and fold changes determined using the ΛΛCT method.

### Tissue fixation, immunohistochemistry and imaging

Third-instar larval and adult tissues were dissected and fixed in 4% formaldehyde in phosphate-buffered saline (PBS) for 30 minutes at room temperature. Tissues were washed in 0.1% Triton-X in PBS (PTX), blocked in PTX with 5% normal goat serum (Amersham Biosciences) for 1 hour at room temperature, and then incubated with primary antibodies overnight at 4°C in block solution. Primary antibodies used were: anti-Prospero (1:20), anti-Repo (1:300), anti-FasIII (1:200), anti-LamC (1:100), and anti-Vasa (1:300) from the Developmental Studies Hybridoma Bank (DSHB), and anti-GFP (1:500, A6455, Thermo Fisher Scientific). Tissues were washed three times for 15 minutes in PTX followed by incubation with secondary antibodies (1:500, anti-mouse or anti-rabbit Alexa 488, Thermo Fisher Scientific) for 1.5 hours at room temperature followed by further washes in PTX. Tissues were stained with 4’-6-diamidino-2-phenylindole (DAPI, 1:1000 in H_2_O for 5 min, Merck) and phalloidin (1:1000 in PTX for 20 min, Thermo Fisher Scientific) during the final wash steps before placing on slides in Vectashield mounting media (Vectorlabs). Where antibody staining was not required, fixed tissues were DAPI stained, washed and mounted as described. Images were acquired on a spinning-disk confocal microscope (Olympus CV1000) and presented as maximum intensity projections with individual Z-slices acquired at not more than 1 μm intervals.

### Survival, developmental timing, and ovariole number

Heterozygous, balanced *beba^T2A^*^-*GAL4*^ adults were mated and allowed to lay on apple juice agar plates for 4 hours at a time over two days. At hatching (24 hours later), first-instar *beba^T2A^*^-GAL4^ homozygotes or *beba^T2A^*^-GAL4^/*beba^df^* transheterozygotes were selected by the absence of GFP (balancer chromosome) using a fluorescent stereo microscope (Leica M165FC). Twenty larvae were placed into 5 vials per genotype. For developmental timing measurements, the number of pupae was scored every 8 h and the mean time to pupariation was calculated for each vial. Larvae heterozygous for *beba^T2A^*^-GAL4^ or *beba^df^* alleles, and a wild-type allele were used as controls. Larval-to-pupal viability was calculated as the proportion of first-instar larvae that reached pupariation. For ovariole numbers, ovary pairs were dissected from at least 9 adult females aged 5-7 days post-eclosion per genotype and fixed in 4% formaldehyde in phosphate-buffered saline (PBS) for 30 min at room temperature. Following gentle washing in PTX, ovarioles were separated and counted under a dissection microscope, and the mean of each ovary pair was determined per individual.

### Digestive system polyploidy

Each digestive tract was segmented into six anatomical regions, and a composite Z-stack image was acquired from each region. DAPI channels were extracted from the Z-stacks and analysed for nuclei number and fluorescence intensity using custom Python code (v3.13; https://github.com/johnsonflygroup/beanbag-RTK/tree/main/analysis/polyploidy) in combination with ilastik (v1.4.1.post1-gpu; Berg et al. 2019). Statistical analyses were conducted in R (v4.5.1). Mean nuclear DAPI intensity was compared across genotypes and regions using one- or two-way ANOVA followed by Tukey’s HSD post hoc tests, as appropriate. Assumptions of normality, homogeneity of variance, and ANOVA validity were assessed using Shapiro–Wilk and Levene’s tests. Variables that failed these assumptions were analysed using non-parametric tests (Kruskal–Wallis with Wilcoxon pairwise comparisons).

### Testis hub cell quantification

Composite Z-stacks were separated into individual channels, and ilastik (v1.4.1.post1-gpu; Berg et al. 2019) was used to generate probability maps for hub cell regions (anti-FasIII) and nuclei (DAPI). Probability maps were then processed with custom Python code (v3.13, https://github.com/johnsonflygroup/beanbag-RTK/tree/main/analysis/hub-cells), which applied thresholding, overlaid DAPI and FasIII signals, and extracted hub nuclei number and FasIII-positive hub volume. Statistical analyses were performed in R (v4.5.1). Genotype differences were tested using one-way ANOVA with Tukey’s HSD post hoc comparisons, following verification of assumptions of normality, homoscedasticity, and ANOVA validity (Shapiro–Wilk and Levene’s tests).

### Bioinformatics

The predicted protein sequence encoded by *D. melanogaster beba* (FlyBase release 6.33) was analysed by SignalP (v4.0; Petersen et al. 2011) and TMpred (Hofmann and Stoffel 1993) for signal peptide sequence and transmembrane domain prediction, respectively. N-linked glycosylation sites were predicted using NetNGlyc1.0 (Gupta and Brunak 2002). Protein accessions for human, *Mus musculus* and *Drosophila melanogaster* RTKs and members of the receptor serine-threonine kinase family (for use as an outgroup) were obtained from the National Centre for Biotechnology Information (NCBI). Arthropod RTK sequences for Ret, Tor and Beba homologs were identified with Basic Local Alignment Search Tool (BLAST; Altschul et al., 1990) using *Drosophila melanogaster* query sequences. The expected value threshold was set at 1.0 e-50 for all searches. Candidate Beba homologues were defined as reciprocal best or high-confidence BLAST hits to *Drosophila* Beba that did not cluster with Tor or Ret in subsequent phylogenetic analysis. For reference, all the accessions used are provided in **Supplementary File 1**. The tyrosine kinase domain (TKD) of all RTK sequences were identified using HMMER biosequence analysis (https://www.ebi.ac.uk/Tools/hmmer/; Eddy 2011) and trimmed manually in MEGA 7 (http://www.megasoftware.net/; Kumar et al. 2016). TKD sequences were aligned using ClustalX2 (http://www.clustal.org/clustal2/; Larkin et al. 2007) using default parameters and Bayesian phylogenies generated in MrBayes v3.2.6 (Ronquist et al. 2012). The analysis assumed a gamma-distribution of rates and used a fixed model. The Monte Carlo Markov Chain search was run with four chains for 700,000 generations with trees sampled every 1000 generations. The first 25% of trees were discarded as ‘burn-in’. The standard deviation of split frequencies (convergence diagnostic) reached 0.05 for the human, mouse and fly sequence analysis, and 0.008 for the arthropod analysis. Trees were visualised and edited in FigTree (http://tree.bio.ed.ac.uk/software/figtree/). The human/mouse/*Drosophila* RTK tree was rooted using receptor serine/threonine kinases as an outgroup; the arthropod Ret/Tor/Beba tree was midpoint-rooted.

## Results and Discussion

### *beba* (CG3277) encodes a receptor tyrosine kinase

The *Drosophila* genome annotation used here predicts *beba* to encode a protein of 789 amino acids (Gramates et al. 2022). This sequence comprises a predicted N-terminal signal sequence, a centrally located transmembrane domain (TM), a conserved C-terminal tyrosine kinase domain (TKD) including an intact active site, and several putative N-linked glycosylation sites in the extracellular domain; all features consistent with the architecture of a typical receptor tyrosine kinase (**Figure 1A**). To test whether Beba can signal like an RTK *in vivo*, we overexpressed the entire coding sequence (UAS-*beba*) under the control of several GAL4 drivers. The ubiquitous driver *actin*-GAL4 caused lethality, as did *engrailed* (*en*)-GAL4 (first instar arrest), which expresses in the posterior compartment of each parasegment. *Patched (ptc)*-GAL4, which expresses in a stripe along the centre of the wing imaginal disc between the presumptive veins L3 and L4, produced viable adults with L3-4 wing vein fusion anterior to the anterior crossvein (**Figure 1B**). This phenotype closely resembles ectopic expression of EGFR using *ptc*-GAL4 (**Figure 1B**, Mohler et al. 2000), supporting the idea that Beba can engage RTK-like signalling *in vivo*. Overexpression in the photoreceptor cells of the eye using GMR-GAL4 produced a strong rough eye phenotype (**Figure 1C**), again similar to that seen with EGFR overexpression under GMR control (Basler and Hafen 1988; Simon et al. 1991; Shilo 2003). Using head extracts from these flies, we checked for activation of Akt and ERK kinases known to act downstream of RTKs, typically via the PI3K and Ras/MAPK pathways, respectively. Both phosphorylated Akt (pAkt) and diphosphorylated ERK (dpERK) were upregulated, with a stronger increase in pAkt than dpERK (**Figure 1D, E**). Taken together, the predicted protein architecture, gain-of-function phenotypes and activation of Akt and ERK phosphorylation support the classification of Beba as a functional *Drosophila* RTK.

**Figure 1.**
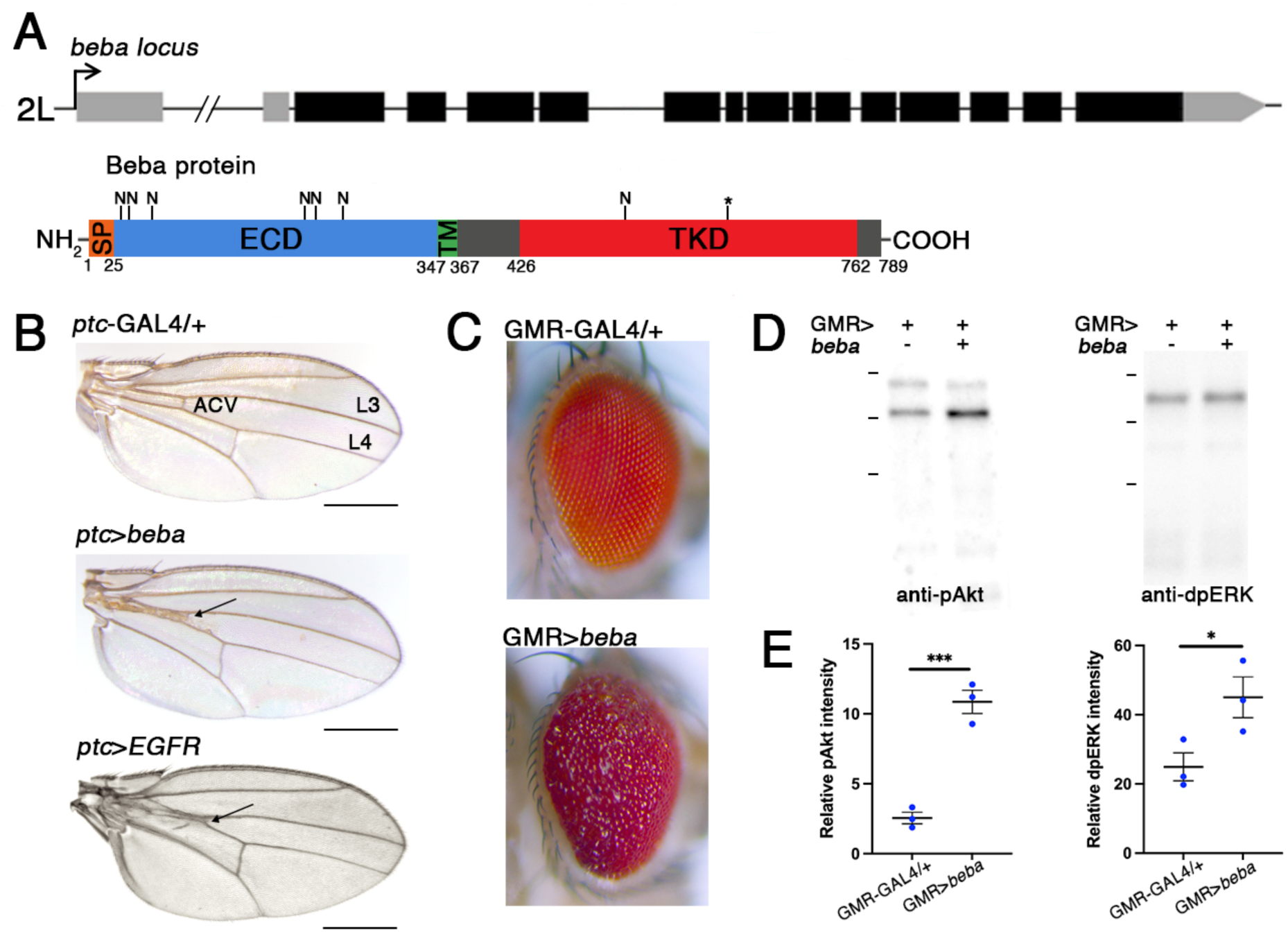
*beba* encodes a receptor tyrosine kinase. (**A**) Gene model of the *beba* locus (untranslated region, light grey; coding sequence, black) and the predicted protein domain architecture, N-linked glycosylation sites (N) and tyrosine kinase active site (asterisk). SP, signal peptide. ECD, extracellular domain. TMD, transmembrane domain. TKD, tyrosine kinase domain. (**B**) Overexpression of *beba* in the patched domain of the wing (*ptc*-GAL4>UAS-*beba*) causes fusion of L3 and L4 anterior of the anterior cross-vein (ACV, arrow). (**C**) Overexpression of *beba* in the eye (GMR-GAL4>UAS-*beba*) produced a strong rough eye phenotype. Immunoblots of head extracts from these flies (**D**) and their quantification (**E**) showed increased levels of the phosphorylated (active) forms of Akt and ERK kinases consistent with activation of PI3K and RAS/MAPK signalling, respectively. *p<0.05, ***p<0.001.

### *Drosophila beba* is expressed in the digestive, nervous and reproductive systems

To better understand the biological roles of *beba*, we next asked which cells and tissues express the gene. For this we made use of a Minos-mediated integration cassette (MiMIC) transposon inserted in the intron separating the fourth and fifth coding exons corresponding to a region N-terminal of the TM domain (**Figure 2A**; Venken et al. 2011). This cassette was replaced with a disruptive Trojan exon containing the T2A peptide followed by GAL4 (Diao et al. 2015; Lee et al. 2018) therefore predicted to truncate Beba before the transmembrane and kinase domains. This design is expected to produce a strong loss-of-function allele while allowing us to observe the endogenous tissue expression pattern of *beba* when placed *in trans* with a UAS-reporter transgene (*e.g.,* UAS-*RedStinger*). To confirm that the trojan exon disrupted *beba* transcription efficiently, we performed quantitative RT-PCR from homozygous *beba^T2A-GAL4^* larvae. As expected, we found that expression of *beba* was severely reduced (**Figure 2B**). This indicates that the *beba^T2A-GAL4^* allele is a strong transcript-disrupting allele.

**Figure 2.**
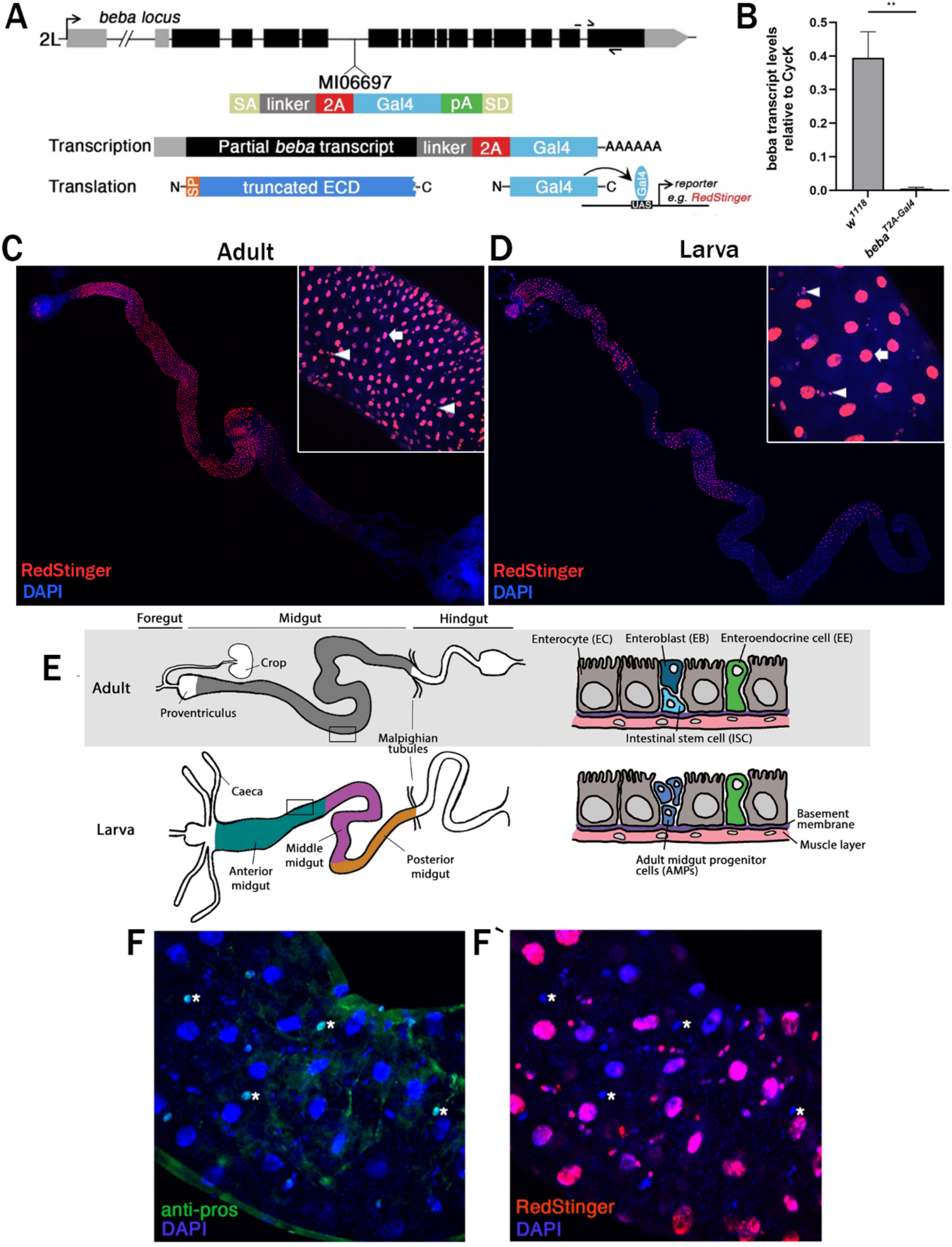
*beba* is expressed in the larval and adult digestive tract. (**A**) The *beba* locus showing position of the MiMIC transposon insertion used (MI06697) and primer locations. The MiMIC cassette was replaced to generate the T2A-GAL4 allele. This is intended to cause premature transcriptional arrest (splice acceptor, SA, and polyadenylation sequence, pA) and translation of truncated beba (partial ECD) and the transcriptional activator GAL4 for use as a reporter. (**B**) Larvae homozygous for the *beba^T2A^*^-*GAL4*^ allele have strongly reduced *beba* expression compared to the wild-type control (p=0.002, n=4). (**C**) Expression of *beba* in the adult digestive tract visualised by UAS-RedStinger expression. Inset: 20x magnification of adult anterior midgut. Arrows denote enterocytes, arrowheads denote smaller cells likely to be intestinal stem cells. (**D**) Expression of *beba* in the larval digestive tract. Inset: 40x magnification of larval anterior midgut. Arrows denote enterocytes, arrowheads denote adult midgut progenitor cells or enteroblasts. (**E**) Schematics of the adult and larval digestive tract showing the structure and regions, cell types, and organisation. (**F**) *beba* is not expressed in larval enteroendocrine cells (anti-pros, green). See nuclei (DAPI, blue) marked by asterisks.

To visualise the *beba* expression pattern, we crossed the *beba^T2A-GAL4^* line to a nuclear-localised fluorescent reporter line (UAS-*RedStinger*) and imaged embryos, larval and adult tissues from the progeny (*beba^T2A-GAL4^*/+; UAS-*RedStinger*). No expression was observed in embryos prior to stage 15. In late-stage embryos expression was weak, whereas in larvae and adults, strong expression was observed in multiple regions of the digestive tract consistent with previous tissue RNA-sequencing surveys (Leader et al. 2018). This included expression in the proventriculus and anterior midgut, and there were also regions with lower or no expression, such as in the posterior midgut and hindgut (**Figure 2C, D**). Higher magnification revealed multiple cell types to be expressing *beba* in the digestive tract. This included the enterocytes (ECs), easily identified because of their large size due to polyploidy (Edgar and Orr-Weaver 2001; Miguel-Aliaga et al 2018), and smaller cells that could be either the enteroendocrine cells (EEs), or the adult midgut progenitor cells (AMPs) in the larva and intestinal stem cells (ISCs) in the adult (**Figure 2E, F;** Micchelli and Perrimon 2006; Ohlstein and Spradling 2006; Miguel-Aliaga et al. 2018). To identify this cell type, we immunostained larval digestive tracts with the EE marker anti-Prospero (anti-Pros, Ohlstein and Spradling 2006). Signal from the RedStinger protein was not observed in the Pros-positive cells, suggesting that the small *beba*-expressing cells in the larval midgut are not EEs and are likely AMPs (**Figure 2G**).

We also detected reporter expression in the larval central nervous system. Confocal imaging revealed expression throughout the entire brain in two populations distinguished by nuclear size and position (**Figure 3A**). We observed large nuclei positioned close to the brain surface across the entire brain, and smaller, often paired nuclei in several regions of the ventral nerve cord (**Figure 3A**). We suspected that the large, likely polyploid nuclei might belong to subperineurial glia (SPG). These are very large flat cells that play a critical role as part of the blood-brain barrier (**Figure 3B**; Schwabe et al. 2005; Unhavaithaya and Orr-Weaver 2012). Consistent with this, we found strong colocalisation with a subpopulation of Repo-positive glia (**Figure 3C**, Halter et al. 1995). This was further confirmed by crossing the *beba^T2A-GAL4^* line to the membrane-bound GFP reporter UAS-mCD8-GFP, which revealed the large membrane outlines that are characteristic of the SPG (**Figure 3D**). This also revealed *beba* expression in a stereotyped set of smaller ventral nerve cord neurons, whose identity remains to be determined (**Figure 3D**).

**Figure 3.**
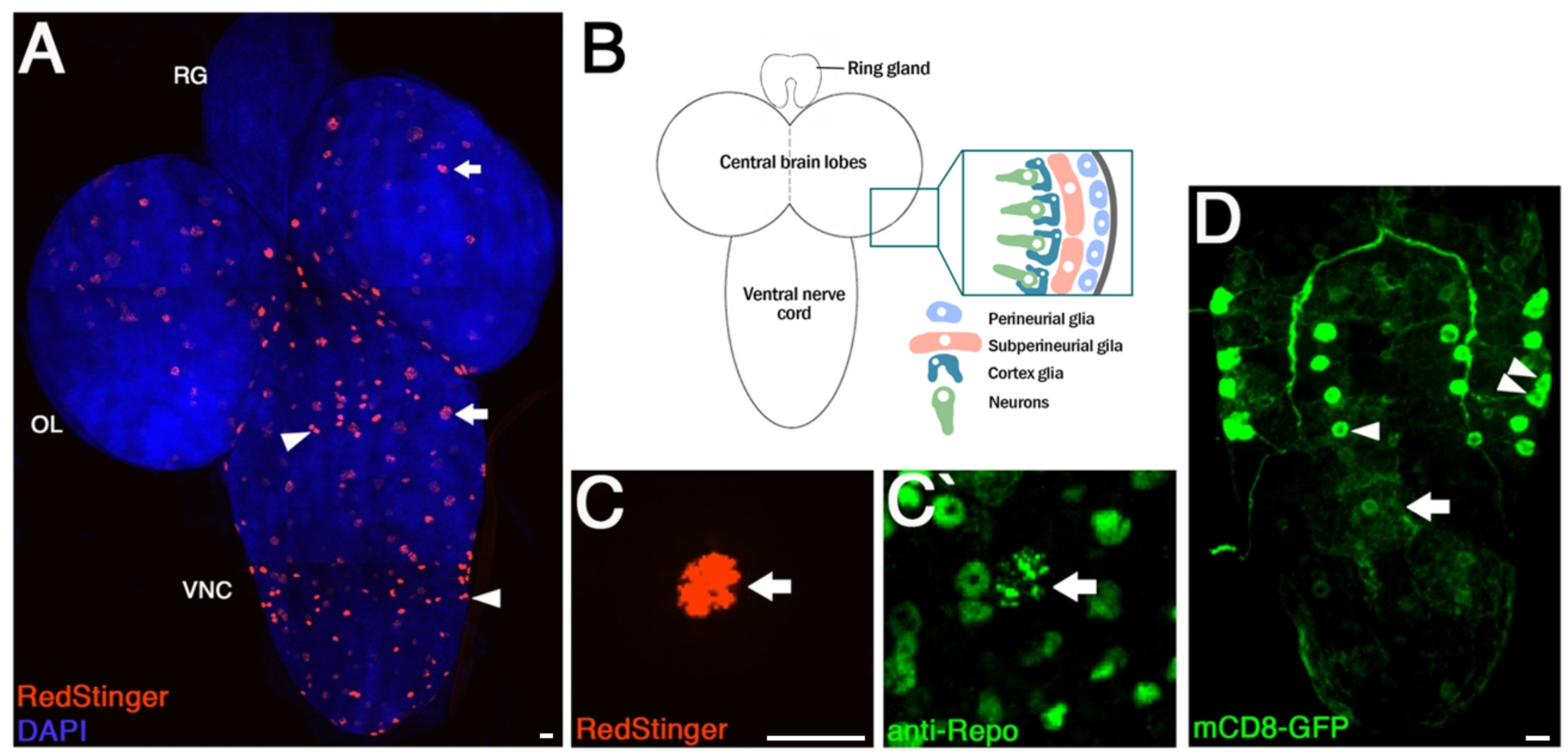
*beba* is expressed in blood-brain barrier glia and a subset of ventral nerve cord neurons. (**A**) Expression of *beba* (driven by UAS-RedStinger) in the larval brain. Expression is observed in large nuclei (arrows) and small, often paired nuclei (arrowhead). RG, ring gland. OL, optic lobe. VNC, ventral nerve cord. (**B**) Schematic of the larval brain structure and cell types of the blood-brain barrier. (**C**) Large nuclei are subperineurial glia (marked by anti-Repo, green, arrowed). (**D**) *beba* expression in the VNC (driven by UAS-mCD8-GFP to mark membranes) reveal neuronal subsets (arrowheads) and large cell boundaries indicative of the subperineurial glia (arrow). Scale bars = 16 µm.

We next dissected the reproductive organs from adults to look for *beba* expression. The testis comprises a spiral-shaped organ with the apical tip housing the germline stem cell niche (**Figure 4A**; Kiger et al. 2001; Tulina and Matunis 2001). Here, expression of *beba^T2A-GAL4^* was limited to a small cluster of cells in the hub region which we confirmed as hub cells by colocalisation with anti-FasIII (**Figure 4B**; Leatherman and DiNardo 2010). In the ovary, we observed highly restricted expression at the anterior tip of the germarium, the region containing the somatic terminal filament and cap cells that organise the female germline stem cell niche (**Figure 4C, D**; Xie and Spradling 2000; Song et al. 2002; Gilboa 2015). LamC marks both terminal filament and cap cells (Gilboa 2015); however, the anterior position of the *beba^T2A^*^-GAL4^-positive cells, together with their overlap with LamC, supports their identity as terminal filament cells (**Figure 4E**). Consistent with this, *beba^T2A^*^-GAL4^ expression was not observed in all LamC-positive cells, suggesting that *beba* expression is restricted to a subset of the LamC-positive niche, most likely the terminal filament. This is consistent with single-cell RNA-sequencing analysis of the developing ovary, in which *beba* expression was reported to be restricted to terminal filament cells (Slaidina et al. 2020). Together, these expression patterns suggest that *beba* may function in somatic reproductive niche cells.

**Figure 4.**
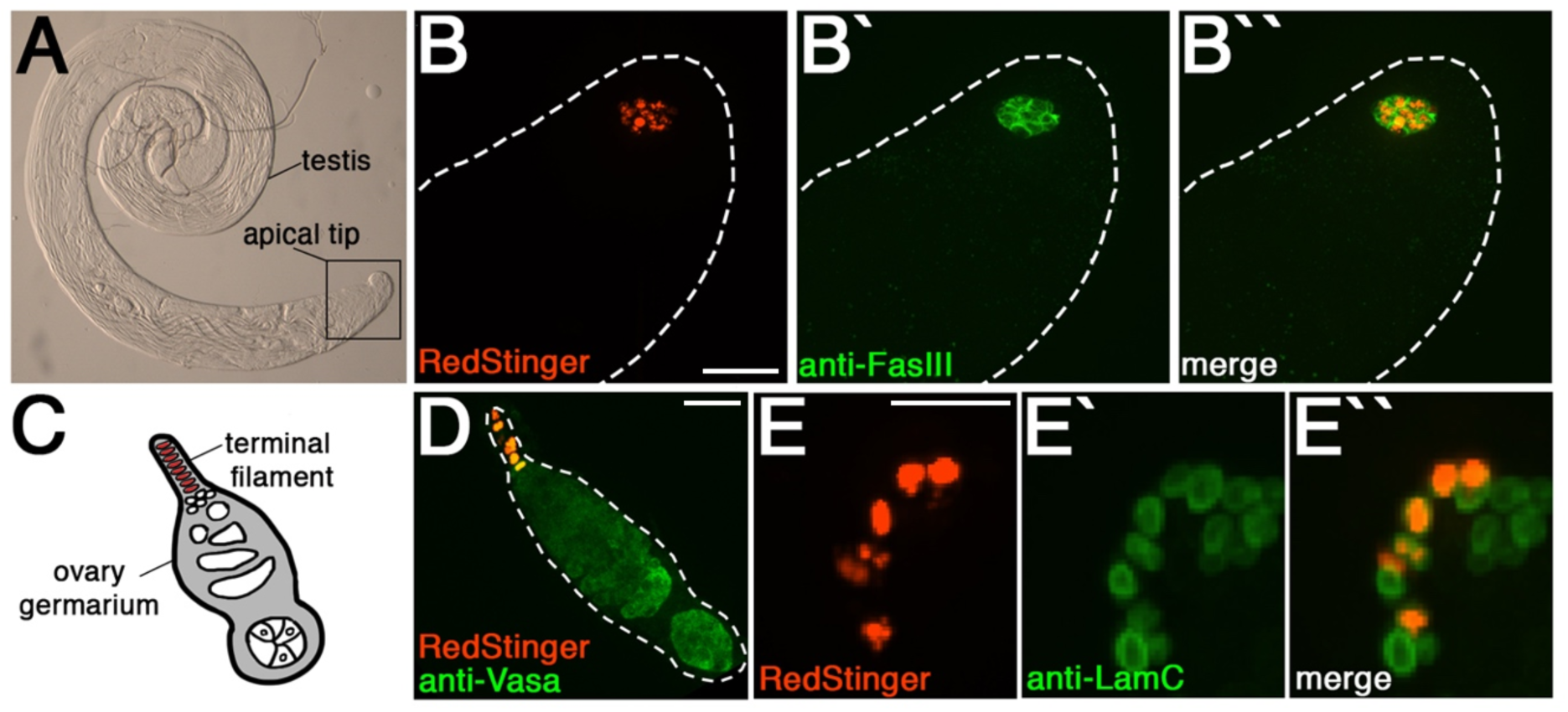
*beba* is expressed in the adult reproductive tissues. (**A**) The adult testis is a spiral-shaped organ with germline stem cells located at the apical tip. (**B**) *beba* expression (*beba^T2A^*^-GAL4^>RedStinger) is found in the hub cell cluster (marked by anti-FasIII, green). (**C**) Schematic of the adult ovary germarium. (**D**) *beba* is expressed (RedStinger) in cells anterior of the germline (marked by anti-Vasa, green). (**E**) Reporter-positive anterior germarial cells are LamC-positive, consistent with terminal filament identity. Scale bars = 16 µm.

From this survey, the expression pattern of *beba* points to two possible biological themes: somatic niche organisation and the support of specialised polyploid cells. In several tissues, *beba* was detected in cells that contribute to tissue architecture or long-term maintenance, including reproductive niche cells and intestinal progenitor-associated populations (Micchelli and Perrimon 2006; Ohlstein and Spradling 2006; Gilboa 2015; Miguel-Aliaga et al. 2018). Its expression in large enterocytes and subperineurial glia also raises the possibility that Beba may participate in pathways linked to endocycle-associated growth or polyploid cell maintenance (Edgar and Orr-Weaver 2001; Unhavaithaya and Orr-Weaver 2012). Given the established roles of RTK-linked pathways such as EGFR/Ras/MAPK and insulin/PI3K signalling in growth, epithelial renewal and tissue homeostasis (Jiang et al. 2011; O’Brien et al. 2011; Miguel-Aliaga et al. 2018), these patterns suggested that *beba* might function in somatic niche regulation or polyploid cell biology.

### *Drosophila beba* is dispensable for viability but required for normal reproductive niche architecture

We next investigated the consequences of *beba* loss using the *beba^T2A-GAL4^* allele. General health was assessed by quantifying viability and the timing of developmental transitions. *beba^T2A-GAL4^*homozygous larvae survived to pupariation in similar numbers to controls, as did transheterozygote individuals for *beba^T2A-GAL4^* and a deficiency allele that includes deletion of *beba (beba^df^,* **Figure 5A**). The homozygotes, however, were significantly delayed in reaching pupariation by more than two days compared to controls (**Figure 5B**), and unlike the transheterozygotes, very few survived until adulthood. Because this delay was not observed in *beba^T2A^*^-GAL4^/*beba^Df^*animals, we interpret the homozygous delay and poor adult recovery as likely arising from background variation or a second-site lesion on the *beba^T2A^*^-GAL4^ chromosome rather than from loss of *beba* itself. Adult transheterozygotes were fertile and showed no obvious external morphological defects. Thus, *beba* is not required for larval-to-adult viability under standard laboratory conditions.

**Figure 5.**
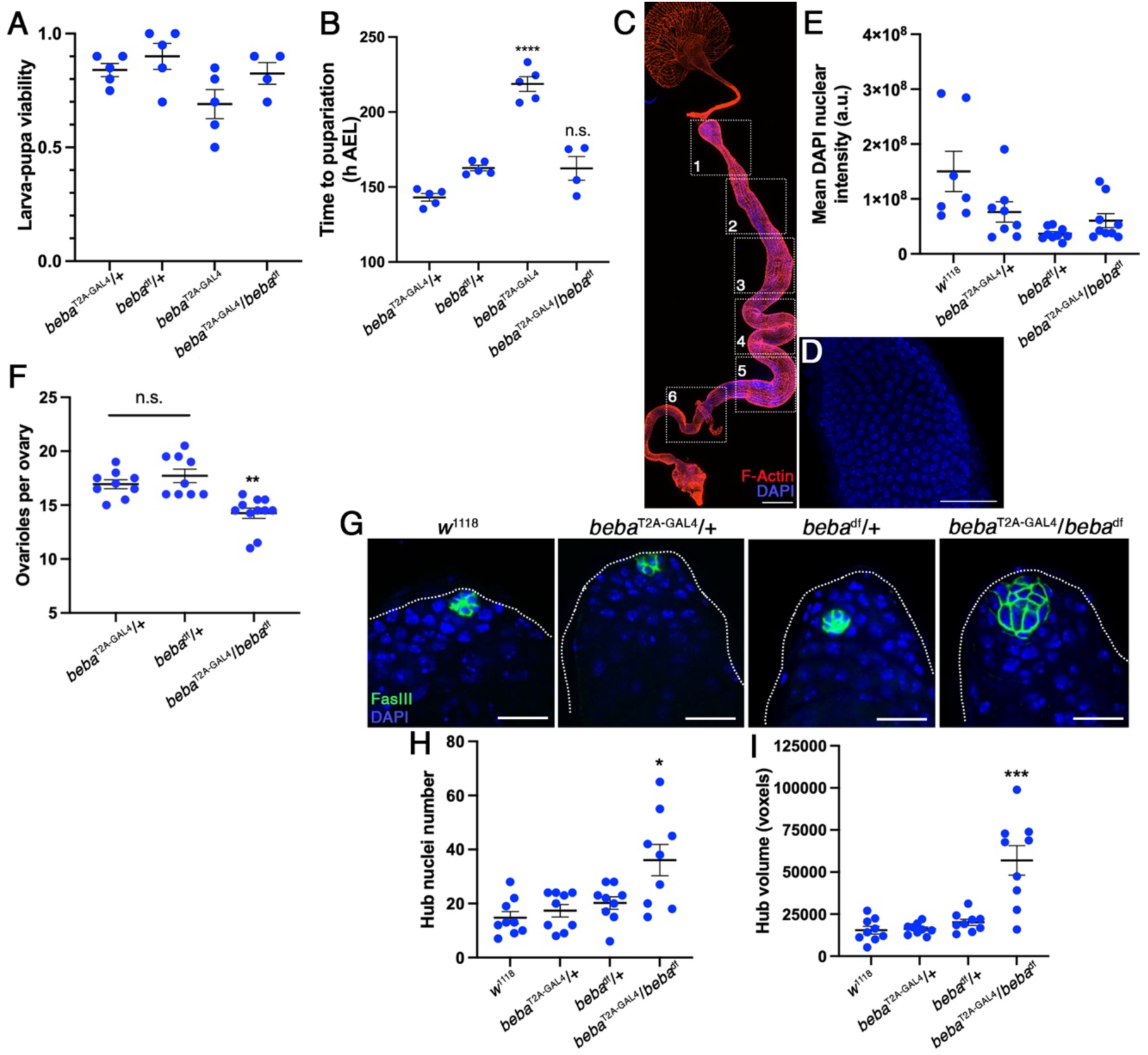
*beba* is dispensable for viability but required for normal female and male reproductive niche architecture. Larval-to-pupal viability (**A**) and developmental timing (**B**) are unaffected in *beba*^T2A-GAL4^/*beba*^Df^ transheterozygotes. *beba^T2A^*^-GAL4^ homozygotes are significantly delayed in development time (p<0.0001). The absence of this delay in *beba^T2A^*^-GAL4^/*beba^Df^* animals suggests that the delay in *beba^T2A^*^-GAL4^ homozygotes is not caused by *beba* loss alone. (**C**) Confocal composite of a representative adult digestive tract showing approximate locations of regions 1-6 used for DNA (DAPI) intensity quantification as a proxy for polyploidy (scale bar is 300 µm). (**D**) Representative example of wild-type (*w*^1118^) enterocyte nuclei (DAPI) from a single region used for quantification (scale bar is 40 µm). (**E**) Mean DAPI intensity per nucleus as a proxy for DNA content showing no effect of *beba^T2A^*^-GAL4^/*beba*^Df^ on enterocyte polyploidy (n=7-9 animals). (**F**) Ovariole number in adult *beba^T2A^*^-GAL4^/*beba*^Df^ females is reduced compared to heterozygote controls (p=0.003, n=9-11 ovary pairs). (**G**) Testis hubs showing nuclei (DAPI) and hub cells (anti-FasIII). Hub cell clusters in *beba^T2A^*^-GAL4^/*beba*^Df^ appeared larger and disorganised (scale bar is 20 µm). Quantification confirmed *beba^T2A^*^-GAL4^/*beba*^Df^ hubs to contain around twice the nuclei on average (**H**, p=0.019, n=9 animals) and a larger cluster volume (**I**, p<0.001, n=9 animals).

We then asked whether *beba* may be supporting endocycle-associated growth or polyploid cell maintenance since its expression is found in the brain subperineurial glia and enterocytes of the digestive tract, both of which are large, highly polyploid cells (Edgar and Orr-Weaver 2001; Unhavaithaya and Orr-Weaver 2012; Miguel-Aliaga et al. 2018). To test this, we investigated the DNA content of polyploid enterocytes in the adult digestive tract using DAPI staining and confocal microscopy. As *beba* did not appear uniformly expressed across the digestive tract, we sampled six regions to capture any potentially local changes to polyploidy (**Figure 5C,D**). We did not detect a significant genotype-dependent change in nuclear DAPI intensity in any region, nor an overall genotype effect when data were pooled across regions (**Figure 5E**). These data do not exclude a context-dependent role for *beba* during epithelial stress, damage or infection, conditions under which RTK signalling is strongly engaged to support intestinal regeneration (Jiang et al. 2011; O’Brien et al. 2011; Miguel-Aliaga et al. 2018). Additionally, since we detect expression in midgut progenitor-like cells, *beba* may instead support progenitor proliferation, differentiation or epithelial repair.

Since *beba* is expressed in reproductive tissues and we observed that *beba^T2A-GAL4^/beba^df^* transheterozygotes retained fertility, we reasoned that loss-of-function phenotypes might be subtle. As *beba* is expressed in the ovarian terminal filament, and this organ has been implicated in regulating final adult ovariole number (Sarikaya et al. 2019; Gilboa 2015), we quantified this trait in *beba^T2A-GAL4^/beba^df^* females. These individuals had a significantly reduced number of ovarioles compared to both heterozygote controls (mean±SEM: *beba^T2A-GAL4^/beba^df^*14.25±0.48; *beba^T2A-GAL4^*/+ 16.94±0.41; *beba^df^*/+ 17.72±0.62; **Figure 5F**). These data suggest that *beba* contributes to normal ovariole formation or terminal filament organisation during ovary development.

We next examined the adult male testis hub, where *beba* is also expressed. Hub cells form a somatic signalling centre that maintains adjacent germline stem cells and cyst stem cells (Kiger et al. 2001; Tulina and Matunis 2001; Leatherman and DiNardo 2010). To investigate *beba* here, we quantified hub cell number and overall hub cluster volume using anti-FasIII to define the hub cluster and DAPI to define nuclei. This revealed significantly elevated hub nuclei number (∼2-fold) and a larger overall hub cluster volume in *beba^T2A-GAL4^/beba^df^*testes compared to both heterozygous and wild-type controls (**Figure 5G-I**). In addition, the FasIII-positive hub domain appeared less compact and more irregularly organised in *beba^T2A-GAL4^*/*beba^Df^* testes (**Figure 5G**). This phenotype could reflect altered hub specification during gonad development, inappropriate proliferation or survival of hub cells, or a defect in maintaining hub architecture after niche formation. Together, the reproductive phenotypes in both females and males indicate that *beba* has non-redundant roles in somatic gonadal niche organisation.

### Beba represents a distinct insect receptor tyrosine kinase family

Beba has not been previously characterised and therefore we were unsure as to its evolutionary origins and whether it was related to other known RTK families. To determine the RTK family to which Beba belongs and whether its conservation extends to mammals, we first performed a Bayesian phylogeny of *Drosophila* Beba together with all known *Drosophila* RTKs and their homologs in mice and humans. This recovered several established RTK family clades with high posterior support (>0.90) on many of the branches, with the receptor serine/threonine kinase outgroup resolved outside the RTK clades (**Figure 6**; Manning et al. 2002; Lemmon and Schlessinger 2010; Mele and Johnson 2020). Beba was found in a moderately supported clade with no unifying mammalian subfamily, alongside *Drosophila* (Dm)Tor, DmRet and mouse Ret, human and mouse Tie, and human Nok. Within this clade there are RTKs that represent different subfamilies, suggesting that these families are closely related and have diverged from a common ancestor.

**Figure 6.**
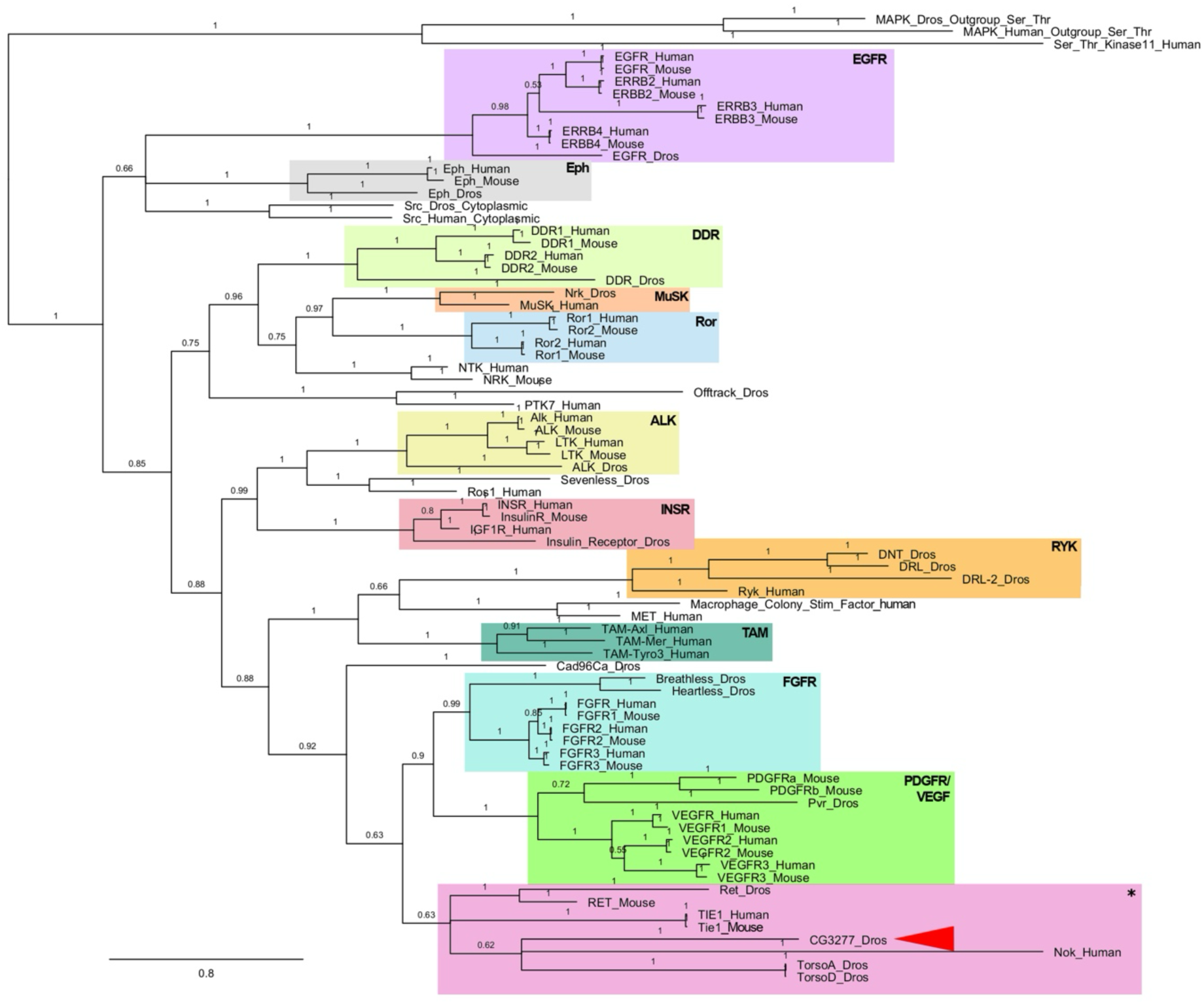
Beba does not cluster within established mammalian RTK families. Bayesian phylogeny of protein tyrosine kinase domains of RTKs present in humans, mice and *Drosophila*. Clades in coloured boxes represent RTKs from the same families that cluster together. Singletons are uncoloured. Posterior probabilities are shown on branches and tree was midpoint rooted using receptor serine/threonine kinases as an outgroup. Beba (CG3277, red arrowhead) is present in a heterogeneous cluster of sequences including Ret, Tie, Nok, and Torso denoted by an asterisk.

Since Beba showed no clear links to any mammalian RTK family, we suspected that it might have been a lineage-restricted RTK that arose within, or was retained primarily within, arthropods. To test this, we gathered Beba homologs from arthropods using BLAST. Initial searches recovered Beba-like candidate sequences from several insect species and more divergent Ret/Tor/Beba-like sequences from non-insect arthropods. Using our search criteria, we did not recover a clear Beba orthologue from *B. mori* or *P. humanus*. With the sequences we found, we performed a second analysis in which Ret and Tor homologs (collected in the same manner) were included from several arthropod species selected to capture the arthropod phylum (for accessions see **Supplementary File 1**). This produced three well-supported clades clearly distinguishing Tor, Ret and Beba homologs, and assigned several of the distant putative Beba-like sequences to the Ret clade (**Figure 7**). Thus, these data suggest that Beba belongs to a distinct insect RTK family related to the Ret/Tor radiation (Manning et al. 2002; Duncan et al. 2014; Mele and Johnson 2020). The absence of confidently assigned Beba orthologues from non-insect arthropods in our analysis is consistent with either an insect-specific origin or retention in insects after loss or extreme divergence in other arthropod lineages.

**Figure 7.**
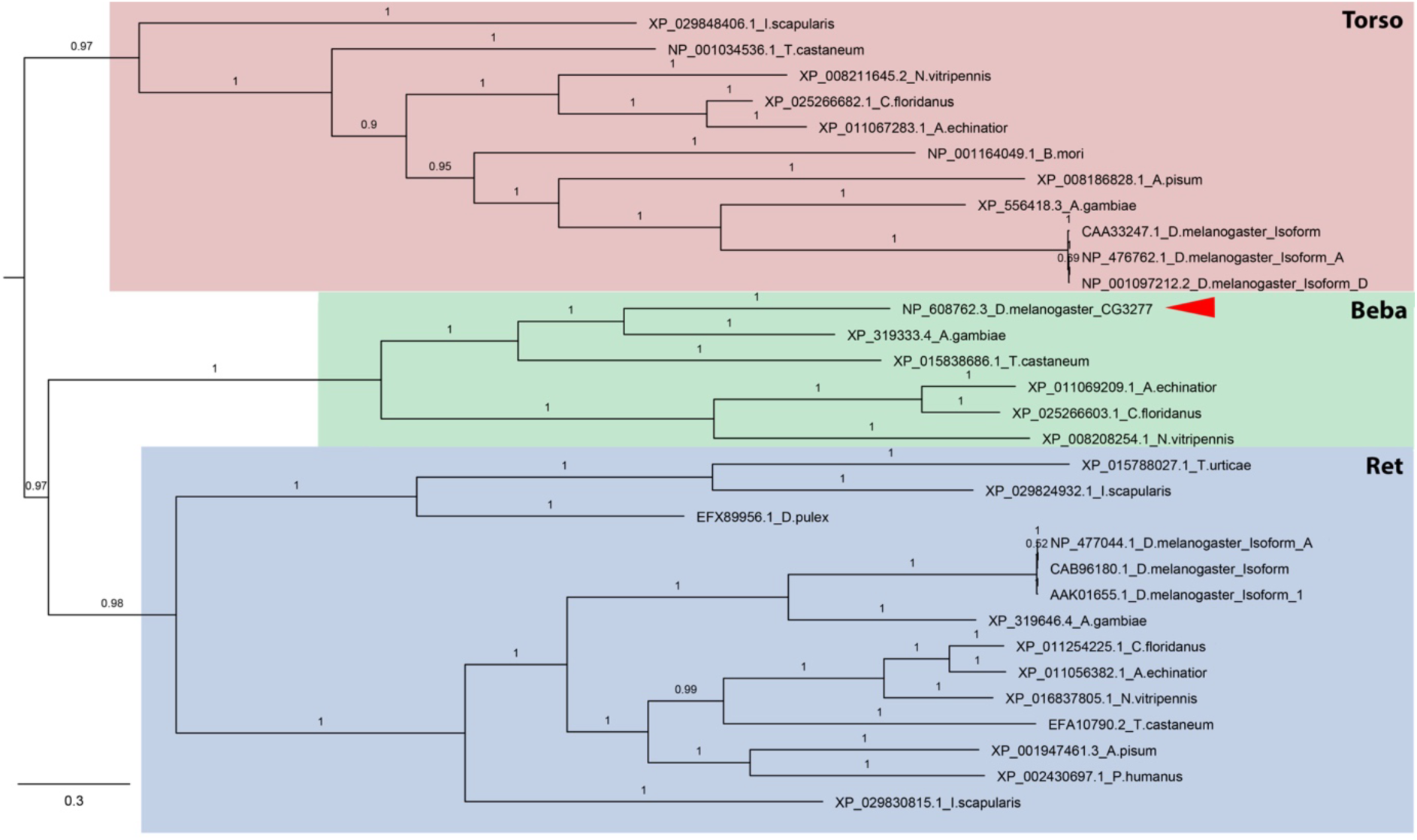
Beba belongs to a distinct insect RTK family. Bayesian phylogeny of arthropod tyrosine kinase domains from Ret, Torso and candidate Beba homologues. The analysis resolves three clades corresponding to Ret, Torso and Beba. Ret (blue) and Torso (pink) clades include representatives from insects, crustaceans and chelicerates, whereas the confidently assigned Beba clade (green) comprises only insect sequences. *Drosophila* Beba (CG3277) is indicated by a red arrowhead. Posterior probabilities are indicated on branches. Species included are *Acromyrmex echinatior, Anopheles gambiae, Acyrthosiphon pisum, Bombyx mori, Camponotus floridanus, Daphnia pulex, Drosophila melanogaster, Drosophila grimshawi, Ixodes scapularis, Nasonia vitripennis, Pediculus humanus, Tribolium castaneum, Tetranychus urticae*.

In summary, we identify *beanbag* as encoding a previously uncharacterised *Drosophila* receptor tyrosine kinase with restricted expression in somatic support cells of the digestive, nervous and reproductive systems. Although *beba* is dispensable for viability under standard laboratory conditions, loss of *beba* alters two reproductive niche-associated structures, reducing ovariole number in females and increasing hub cell number and hub volume in males, indicating a specialised role in somatic gonadal organisation. The ligand that activates Beba remains unknown, but its relationship to the Torso and Ret RTK branches raises the possibility that Beba may use a regulated extracellular ligand or cofactor. In *Drosophila*, Torso is activated by the cysteine-knot growth factor Trunk during terminal patterning and by prothoracicotropic hormone during developmental timing, illustrating how related ligands can activate the same receptor in distinct developmental contexts (Casanova et al. 1995; Casali and Casanova 2001; Rewitz et al. 2009; Duncan et al. 2014). Whether Beba responds to a known Torso-associated ligand, a Ret-associated extracellular factor, or a distinct insect-specific ligand remains an important question for future work. More broadly, these findings expand the repertoire of *Drosophila* RTKs and show how lineage-restricted receptors can evolve specialised roles in tissue architecture.

## Acknowledgements

We thank the Bloomington *Drosophila* Stock Centre, Professors Hugo Bellen (Baylor College of Medicine) and Marc Freeman (Vollum Institute) for fly stocks, and the Developmental Studies Hybridoma Bank for antibodies. We further thank FlyBase for genome curation, the Australian *Drosophila* Biomedical Research Support Facility (OzDros) for assistance in importing fly stocks, and Professors Peter Dearden (University of Otago) and Gary Hime (University of Melbourne) for helpful suggestions regarding the phylogenetics and functional work, respectively. This work was supported by an Australian Research Council (ARC) Discovery Project to T.K.J. (DP220103444). T.K.J. is supported by an ARC Future Fellowship (FT220100023).

## Data availability

All data necessary for confirming the conclusions of the article are present within the article, figures, and tables. Strains and plasmids are available upon request. Code for the analysis of intestinal polyploidy and hub cell clusters are available at: https://github.com/johnsonflygroup/beanbag-RTK

